# Direct and indirect nutritional factors that determine reproductive performance of heifer and primiparous cows

**DOI:** 10.1101/2020.03.20.000315

**Authors:** Lidiane R. Eloy, Carolina Bremm, José F. P. Lobato, Luciana Pötter, Emilio A. Laca

## Abstract

Pregnancy rate is a major determinant of population dynamics of wild ungulates and of productivity of livestock systems. Allocation of feeding resources, including stocking rates, prior to and during the breeding season is a crucial determinant of this vital rate. Thus, quantification of effects and interaction among multiple factors that affect pregnancy rate is essential for management and conservation of pasture-based systems. Pregnancy rate of 2982 heifers and primiparous cows was studied as a function of animal category, average daily gain during the breeding season, stocking rate, pasture type and body weight at the beginning of the breeding season. Data were obtained from 43 experiments conducted in commercial ranches and research stations in the Pampas region between 1976 and 2015. Stocking rate ranged from 200 to 464 kg live weight/ha, which brackets values for most of the grazinglands in similar regions. Age at breeding was 14-36 months (24.6 ± 7.5 months); initial breeding weights were 129-506 kg and 194-570 kg for heifers and primiparous cows. Pregnancy rate was modeled with an apriori set of explanatory variables where proximate variables (breed, body weight at start of breeding, weight gain during breeding and category) were included first and subsequently modeled as functions of other variables (pasture type, supplementation and stocking rate). This modeling approach allowed detection of direct and indirect effects (through nutrition and body weight) of factors that affect pregnancy rate. Taurine (*Bos taurus* breeds, N = 1058) had higher pregnancy rate than *B. Taurus* x *B. indicus* crossbreed (N = 1924) females. Pregnancy rate of heifers and primiparous cows grazing in natural grasslands decreased with increasing stocking rate, but no effect of stocking rate was detected in cultivated and improved pastures. Pregnancy rate increased with increasing average daily gain during the breeding season. Use of cultivated or improved natural pastures promotes higher pregnancy rate, as well as allows higher in stocking rate at the regional level. Body weight at the start of the breeding season is the primary determinant of pregnancy rates in heifer and primiparous cows.

## Introduction

Although fundamental cattle physiology and its relation to nutrition are well known, there is not sufficient quantitative information about how management affects nutrition and reproduction, especially under natural grasslands conditions. In particular, few studies have been done to evaluate the relative effects of stocking rate and other nutritional management factors on pregnancy rates at a regional level (Gottschall & Lobato 1996; Simeone & Lobato 1996; Quadros & Lobato 1996; Fagundes, Lobato & Schenkel 2003; Pötter *et al.* 2010; Sartori & Guardieiro 2010). The lack of single studies with regional scope is understandable because such studies with livestock are logistically complex and extremely expensive.

An alternative to specific comprehensive studies is to analyze data pooled form multiple studies (Duffield, Merril & Bagg 2012; Lean, Thompson & Dunshea 2014) that address the same research question using equivalent response and explanatory variables, that is, a meta-analysis (Schwarzer, Carpenter & Rücker 2015). Meta-analysis requires care in the process of systematizing results from multiple studies but it has the advantage of increasing precision, decreasing costs and research time, and increasing the degrees of freedom in the analysis (Fisher 1999; Lovatto, Lehnen, Andretta, Carvalho & Hauschild 2007).

Livestock production in the Pampas region of Southern Brazil is characterized by a low pregnancy rate that has remained stagnant over many years, despite multiple changes in economic and technological factors that affect productivity (IBGE 2020). Low pregnancy rates prevent full development of the livestock sector in many regions of the world, and it may be associated with poor pasture management, overstocking and lack of differential nutritional management for animal categories with different requirements. Stocking rate is considered the most important decision in grazing management because it affects the forage base, herbage allowance, intake and animal performance (Sollenberger, Agouridis, Vanzant, Franzluebbers & Owens 2012). Nutritional limitation during periods of high requirement can compromise development and delay puberty of heifers, as well as inhibit ovulation of cows (Rocha & Lobato 2002).

Time at which puberty occurs relative to the start of breeding season is what determines pregnancy rate in the first breeding season of heifers (Sparke & Lamond 1968), which influences a cow’s ability to get pregnant in subsequent years and remain in the herd, determining her lifetime productivity. Puberty of heifers is influenced by management of the annual production cycle, as well as the physiology (production and release of hormones) and its genetic (breed and size of mature age) (Day 2015). In addition to the use of pastures, body weight at the beginning of the breeding season is associated with animal nutrition and it is an important factor influencing the reproductive performance of heifers and beef cows (Richards, Spitzer & Warner 1986; Osoro & Wring 1992; Roso *et al*. 2009).

Natural grasslands and cultivated pastures constitute the forage basis for beef cow herds in many regions of the world, including the US (USDA-ERS 2020). The grasslands that support cow-calf operation in the Pampas are characterized by spring-summer growth, with quality and availability reduced in autumn and winter. Cultivated and improved pastures are utilized to satisfy the nutritional requirements of cattle, especially during the cooler months when natural pasture growth is limited. Combined with cultivated and improved pastures, supplements may be used to increase average daily gain of grazing animals and to promote greater reproductive development (Frizzo *et al.* 2003).

The aim of the present study was to integrate information from multiple studies of factors that affect pregnancy rates under production conditions in the Pampas to carefully quantify response curves relating pregnancy rate to the most important predictors. First, we take an approach where pregnancy rate is analyzed as a function of known proximate factors such as body weight, category and changes in body weight per day during the breeding season. Second, we add the effects of stocking rate, supplementation and pasture type on proximate factors and directly on pregnancy rate to account for effects not mediated by the proximate factors evaluated. Because of the long term, large geographic region and large number of cows involved in this synthesis, results should be useful not only for ranch-level management but also for regional agricultural policy.

## Materials and Methods

### Study Sample

Data used in this paper were obtained from published literature. Experiments were conducted at the Agronomic Experimental Station of Federal University of Rio Grande do Sul and in private ranches in Rio Grande do Sul, Southern Brazil, to investigate the effects of several factors on pregnancy rate of heifers and primiparous cows (Table 1) between 1976 and 2015. Data include records from 29 doctoral dissertations or master’s theses for a total of 43 experiments (some studies had more than one experiment).

Experiments were selected because the original raw data were available for all of them. According to Köppen (Wrege, Steinmetz, Reisser Junior & Almeida 2011), climate in all sites represented in the data is subtropical humid.

**Table 1.** Relation of studies of database with n (number of animals), location, coordinated geographic, precipitation and type of pasture in southern Brazil

| Author | n | Local | Coordinated geographic | Precipitation (mm/year) | Type of pasture* |
| --- | --- | --- | --- | --- | --- |
| Albospino, 1990 | 23 | Eldorado do Sul | 30°52'/51°39' | 1332 | Italian ryegrass (*Lolium multiflorum* Lam.)<br>Arrowleaf clover (*Trifolium vesiculosum*) |
| Azambuja, 2003 | 216 | Arambaré | 31°11'/51°74' | - | Natural pasture |
| Beretta, 1994 | 113 | Eldorado do Sul | 30°52'/51°39' | 1398 | Annual ryegrass (*Lolium multiflorum* Lam.)<br>Arrowleaf clover (*Trifolium vesiculosum*) |
| Cachapuz, 1976 | 57 | Dom Pedrito | 30°99'/54°70' | 1376 | Natural pasture<br>Common vetch (*Vicia sativa*) |
| Deresz, 1976 | 110 | Pelotas | 30°58'/50°40' | 1285 | Natural pasture<br>Annual ryegrass (*Lolium multiflorum* Lam.)<br>Arrowleaf clover (*Trifolium vesiculosum*) |
| Fagundes, 2001 | 87 | Itaqui | 29°24'/56°47' | 1500 | Natural pasture |
| Freitas, 2005 | 350 | São Gabriel | 30°33'/54°32' | 1193 | Annual ryegrass (*Lolium multiflorum* Lam.)<br>White clover (*Trifolium repens*)<br>Bird's-foot trefoil (*Lotus corniculatus*) |
| Gottschall, 1994 | 114 | São Gabriel | 30°33'/54°32' | 1512 | Natural pasture |
| Lopes, 2004 | 39 | Eldorado do Sul | 30°52'/51°39' | 1446 | Black oats (*Avena strigosa*)<br>Annual ryegrass (*Lolium multiflorum* Lam.)<br>Arrowleaf clover (*Trifolium vesiculosum*)<br>Pearl millet (*Pennisetum americanum*) |

\*Natural pasture with prevalence of Bahiagrass (*Paspalum notatum*), Dallisgrass (*Paspalum dilatatum*), *Axonopus affinis*, *Andropogon lateralis*, *Trifolium polymorphum*

Table 1 ... Continued
| Author | n | Local | Coordinated<br>geographic | Precipitation<br>(mm/year) | Type of pasture* |
| --- | --- | --- | --- | --- | --- |
| Magalhães, 1992 | 210 | Rosário do Sul | 30°25'/54°92' | 1550 | Annual ryegrass (*Lolium multiflorum* Lam.) |
| Marques, 2001 | 231 | Eldorado do Sul | 30°52'/51°39' | 1440 | Black oats (*Avena strigosa*)<br>Annual ryegrass (*Lolium multiflorum* Lam.)<br>Arrowleaf clover (*Trifolium vesiculosum*) |
| Menegaz, 2006 | 323 | Uruguaiana | 29°76'/57°09' | 1500 | Natural pasture<br>Annual ryegrass (*Lolium multiflorum* Lam.)<br>White clover (*Trifolium repens*)<br>Bird's-foot trefoil (*Lotus corniculatus*) |
| Moraes, 1991 | 60 | Dom Pedrito | 30°99'/54°70' | 1300 | Annual ryegrass (*Lolium multiflorum* Lam.)<br>White clover (*Trifolium repens*)<br>Sorghum (*Sorghum bicolor*)<br>Bird's-foot trefoil (*Lotus corniculatus*) |
| Müller, 1998 | 50 | Eldorado do Sul | 30°52'/51°39' | 1440 | Annual ryegrass (*Lolium multiflorum* Lam.)<br>Arrowleaf clover (*Trifolium vesiculosum*) |
| Nardon, 1985 | 65 | Eldorado do Sul | 30°52'/51°39' | 1398 | Annual ryegrass (*Lolium multiflorum* Lam.)<br>Arrowleaf clover (*Trifolium vesiculosum*) |
| Pereira Neto, 1996 | 62 | Eldorado do Sul | 30°52'/51°39' | 1332 | Annual ryegrass (*Lolium multiflorum* Lam.)<br>Arrowleaf clover (*Trifolium vesiculosum*) |
| Pilau, 2007 | 234 | Tupanciretã | 29°03'/53°48' | - | Black oats (*Avena strigosa*)<br>Annual ryegrass (*Lolium multiflorum* Lam.) |
| Polli, 1986 | 71 | Eldorado do Sul | 30°52'/51°39' | 1398 | Natural pasture<br>Annual ryegrass (*Lolium multiflorum* Lam.)<br>Arrowleaf clover (*Trifolium vesiculosum*) |

\*Natural pasture with prevalence of Bahiagrass (*Paspalum notatum*), Dallisgrass (*Paspalum dilatatum*), *Axonopus affinis*, *Andropogon lateralis*, *Trifolium* *polymorphum*

**Table 1... Continued**
| Author | n | Local | Coordinated geographic | Precipitation (mm/year) | Type of pasture* |
| --- | --- | --- | --- | --- | --- |
| Pötter, 2002 | 92 | Quaraí | 30°26'/56°01' | 1356 | Natural pasture<br>Annual ryegrass (*Lolium multiflorum* Lam.)<br>Bird's-foot trefoil (*Lotus corniculatus*) |
| Quadros, 1991 | 69 | Dom Pedrito | 30°99'/54°70' | 1540 | Natural pasture |
| Ribeiro, 1986 | 70 | Cachoeira do Sul | 30°03'/52°89' | 1621 | Natural pasture |
| Rocha, 1997 | 394 | Dom Pedrito | 30°44'/54°47' | 1450 | Annual ryegrass (*Lolium multiflorum* Lam.)<br>White clover (*Trifolium repens*)<br>Red clover (*Trifolium pratensis*)<br>Bird's-foot trefoil (*Lotus corniculatus*) |
| Rosa, 2010 | 241 | Dom Pedrito | 30°44'/54°47' | 1300 | Natural pasture<br>Annual ryegrass (*Lolium multiflorum* Lam.)<br>White clover (*Trifolium repens*)<br>Bird's-foot trefoil (*Lotus corniculatus*) |
| Silva, 2010 | 142 | Bagé | 31°22'/54°39' | 1300 | Annual ryegrass (*Lolium multiflorum* Lam.) |
| Simeone, 1995 | 119 | Bagé | 31°22'/54°39' | 1350 | Natural pasture |
| Souza, 2005 | 64 | Dom Pedrito | 30°99'/54°70' | 1376 | Annual ryegrass (*Lolium multiflorum* Lam.)<br>White clover (*Trifolium repens* L.) |
| Souza, 2015 | 49 | Júlio de Castilhos | 29°23'/53°68' | - | Black oats (*Avena strigosa* Schreb.)<br>Palisade grass (*Urochloa brizantha*) |
| Tanure, 2008 | 194 | Quaraí | 30°26'/56°01' | 1356 | Natural pasture |
| Zanotta Jr, 1984 | 84 | Pelotas | 30°58'/50°40' | 1285 | Natural pasture<br>Annual ryegrass (*Lolium multiflorum* Lam.)<br>White clover (*Trifolium repens*) |

\*Natural pasture with prevalence of Bahiagrass (*Paspalum notatum*), Dallisgrass (*Paspalum dilatatum*), *Axonopus affinis*, *Andropogon lateralis*, *Trifolium* *polymorphum*

### Data

The initial database created contained the following variables for each of 3933 animals: age at the beginning of the breeding season was 14 to 36 months (average was 24.6 ± 7.5 months); animal categories were heifers (n = 2257) and primiparous cows (n = 1676); average animal weight was 315.4 ± 55.9 kg at the beginning and 337.3 ± 53.4 kg at the end of the breeding season; body condition score in a 0-5 scale was 3.2 ± 0.6 at the beginning and 3.4 ± 0.6 at the end of the breeding season; breeds were Angus (n = 306), Braford (n = 499), Brangus (n = 323), Crossbred (n = 1928), Devon (n = 110) and Hereford (n = 767); stocking rate ranged from 200 to 464 kg of body weight per hectare (kg BW/ha) and averaged 337.32 ± 54.68 kg BW/ha; pasture types were cultivated (n=2050), improved (n=324) or natural grassland (n=1559); feed supplementation before the breeding season was recorded as a binary variable (supplemented vs. not supplemented).

Breeds were recoded as crossbred (final N = 1924) vs. *B. Taurus* (final N = 1058). One experiment with an extreme stocking rate of 800.0 kg of body weight per hectare was excluded from the analysis. The remaining data had stocking rates ranging from 200 to 463.5 (average was 336.85 ± 54.92) kg BW/ha, which are more typical for the region. Columns with large number of missing values and rows without complete multivariate records were excluded, resulting in a final data set of 2982 records (animals) for which all variables depicted in Fig 1 were available.

**Fig. 1.**
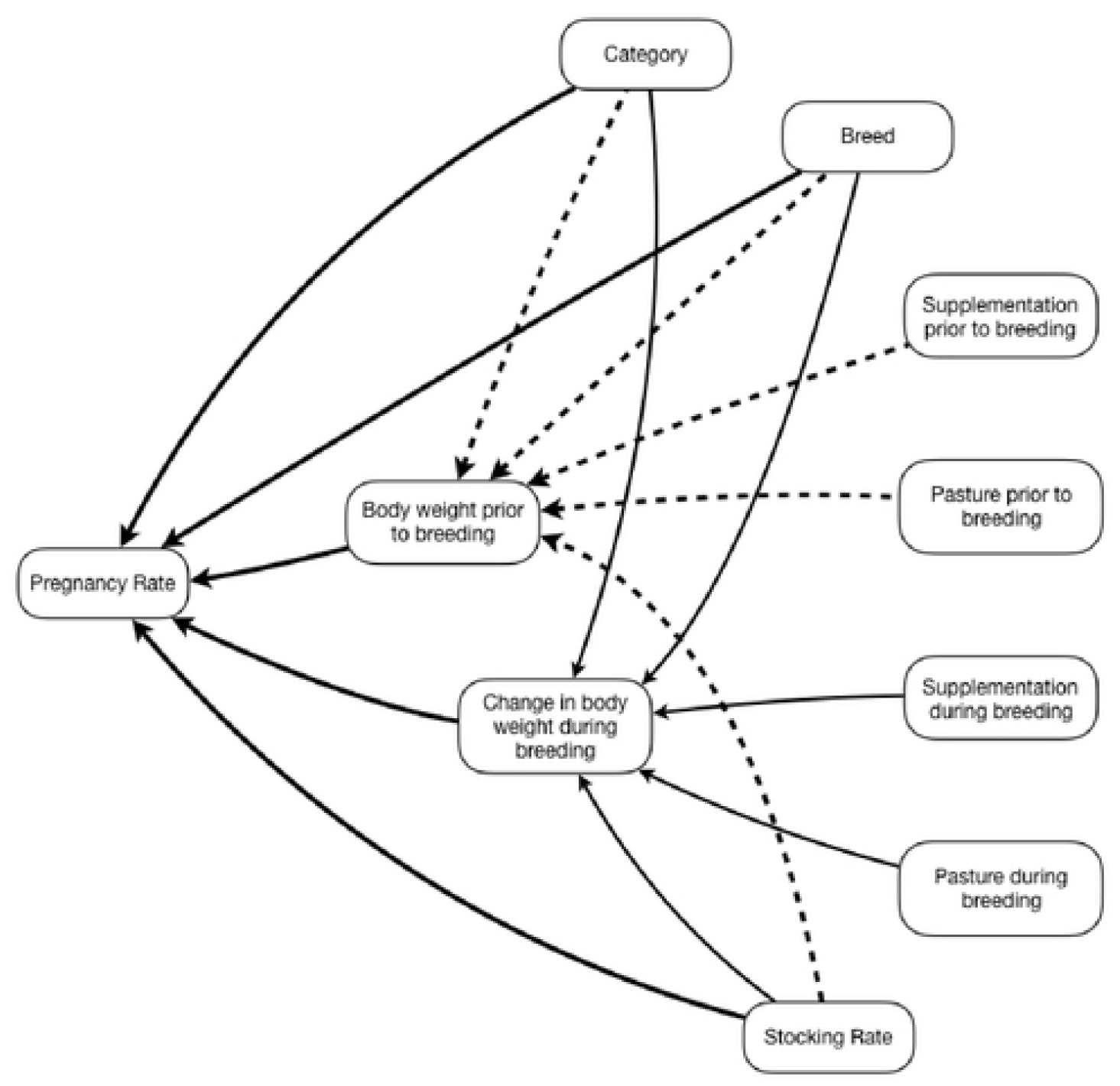
Schematic structure of the hypothesized determinants of pregnancy rate (PR).

### Statistical Analyses

Statistical analyses were conducted using R (R Development Core Team 2015). We used generalized linear mixed models (GLMM) based on the logit link function, as they are generally recommended for binary data (Agresti 2002; Sauvant, Schmidely, Daudin & St-Pierre 2008). Data for each heifer and primiparous cow were available, allowing an analysis analogous to an incomplete block design, with a random intercept for each experiment (Senn, Gavini, Magrez & Scheen 2011).

The main response variable was pregnancy rate as evaluated by the relation between the number of pregnant heifers or primiparous cows and the total number of heifers or primiparous cows in each experiment. Explanatory factors considered were category, weight at the beginning of the breeding season, weight change during the breeding season, breed, stocking rate, and type of pasture before and after the breeding season. All quantitative variables were standardized to facilitate the convergence of the computations to estimate parameters. A structure of causal effects was established a priori (Fig 1) and then simplified by removing nonsignificant components.

First, pregnancy rate (PR) was analyzed using *glmer* with a binomial distribution and a logit link, as a function of known proximate factors such as body weight and category. Breed was included to account for inherent differences in breeds that could modulate the effects of body weight, for example, due to differences in mature body or frame size. Model selection proceeded by simplification of a full model until it had only those effects that were significant or part of significant interactions. Significance of terms was assessed by type II Wald tests using the Anova() function of the *car* package (Fox & Weisberg 2011). The initial full model, expressed as an R formula for the generalized linear mixed-effects model (glmer) function of the lme4 package (Bates, Maechler, Bolker & Walker 2014) was

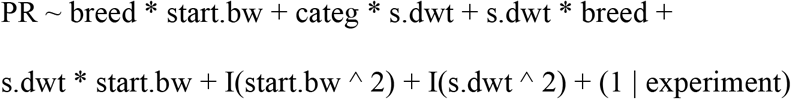

where *categ* is animal category, *start.bw* is weight at the beginning of the breeding season, *s.dwt* is daily weight gain, *I(start.bw^2)* is squared weight at the beginning of the breeding season, *I(s.dwt^2)* is daily weight gain squared and *experiment* is a categorical variable or factor with a different value for each experiment. Each experiment was allowed a random effect to account for the potential intraclass correlation caused by common condition for all animals in each experiment. The “*” operator indicates that both main effects and their interaction are included in the model. This final model was tested against the full model by a likelihood-ratio test using the anova() function to make sure they were not significantly different.

Second, stocking rate was added as the last term to the resulting model to determine if stocking rate had significant effects beyond those effected via proximate factors. Significance of stocking rate effects was assessed with the same Wald test as before. Third, body weight at the beginning of the breeding season was modeled with the following full model:

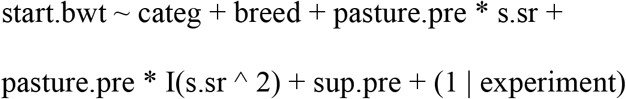

where pasture.pre is a factor indicating whether animals grazed natural grassland, cultivated pastures or improved grassland prior to the breeding period; s.sr is stocking rate, and I(s.sr^2) is the quadratic effect of stocking rate. Other terms were defined above. Finally, change in body weight during the breeding period (s.dwt) was analyzed starting with the following full model:

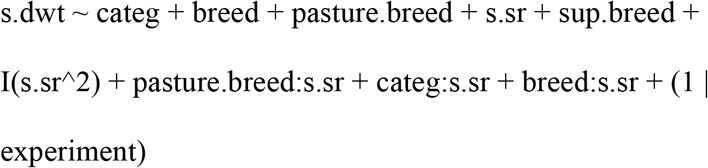

where pasture.breed is the type of pasture grazed during the breeding period and sup.pre is a binary variable indicating whether animals received supplementation during the breeding period. Models for start.bwt and s.dwt were simplified and final models were tested following the same procedures as before. For all models, assumptions were assessed by inspection of residual plots.

## Results

### Effects of proximate causal factors on pregnancy rate

The final model for selected pregnancy rate was:

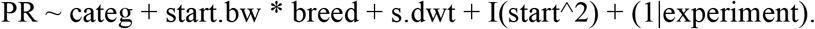

The most important factor affecting pregnancy rate was body weight at the beginning of the breeding season (Table 2), which accounted for 68 % of the model sum of squares. Mc Fadden’s pseudo R^2^ (Domencich & McFadden 1974) for fixed effects of the complete final model was 9.53%, whereas a model with only the linear and quadratic effects of initial body weight had a pseudo R^2^ equal to 8.8%.

**Table 2.** Analysis of variance of the model for pregnancy rate with proximate factors.

| Effect | Wald's Chi-sq (type II) | df | p-value |
| --- | --- | --- | --- |
| start.bw<sup>1</sup> | 234.9 | 1 | <0.0001 |
| I(start.bw<sup>2</sup>)<sup>2</sup> | 48.2 | 1 | <0.0001 |
| s.dwt<sup>3</sup> | 11.1 | 1 | 0.0009 |
| categ<sup>4</sup> | 10.7 | 1 | 0.0011 |
| start.bw:breed<sup>5</sup> | 6.2 | 1 | 0.0141 |
| breed<sup>6</sup> | 2.7 | 1 | 0.0995 |

<sup>1</sup>Weight at the beginning of the breeding period (standardized using mean = 318 kg, s = 69 kg).
<sup>2</sup>Quadratic effect of weight at the beginning of the breeding season.
<sup>3</sup>Average daily gain during the breeding season (standardized using mean = 0.264 kg/day, s = 0.314 kg/day).
<sup>4</sup>Animal category (heifer or primiparous cow).
<sup>5</sup>Interaction between weight at the beginning of the breeding season and breed.
<sup>6</sup> Breed type (crossbred or taurine).

Pregnancy rate increased steeply with increasing body weight at the beginning of the breeding season for crossbreed and taurine females, but for taurine females it increased faster and reached a higher maximum than for crossbreed females. Taurine females starting the breeding season with an average weight of 440.0 kg or more and had an expected pregnancy rate of 99.0 %, whereas crossbred females that started the breeding season with similar weight had an expected pregnancy rate of 91.0 % (Fig 2).

**Fig. 2.**
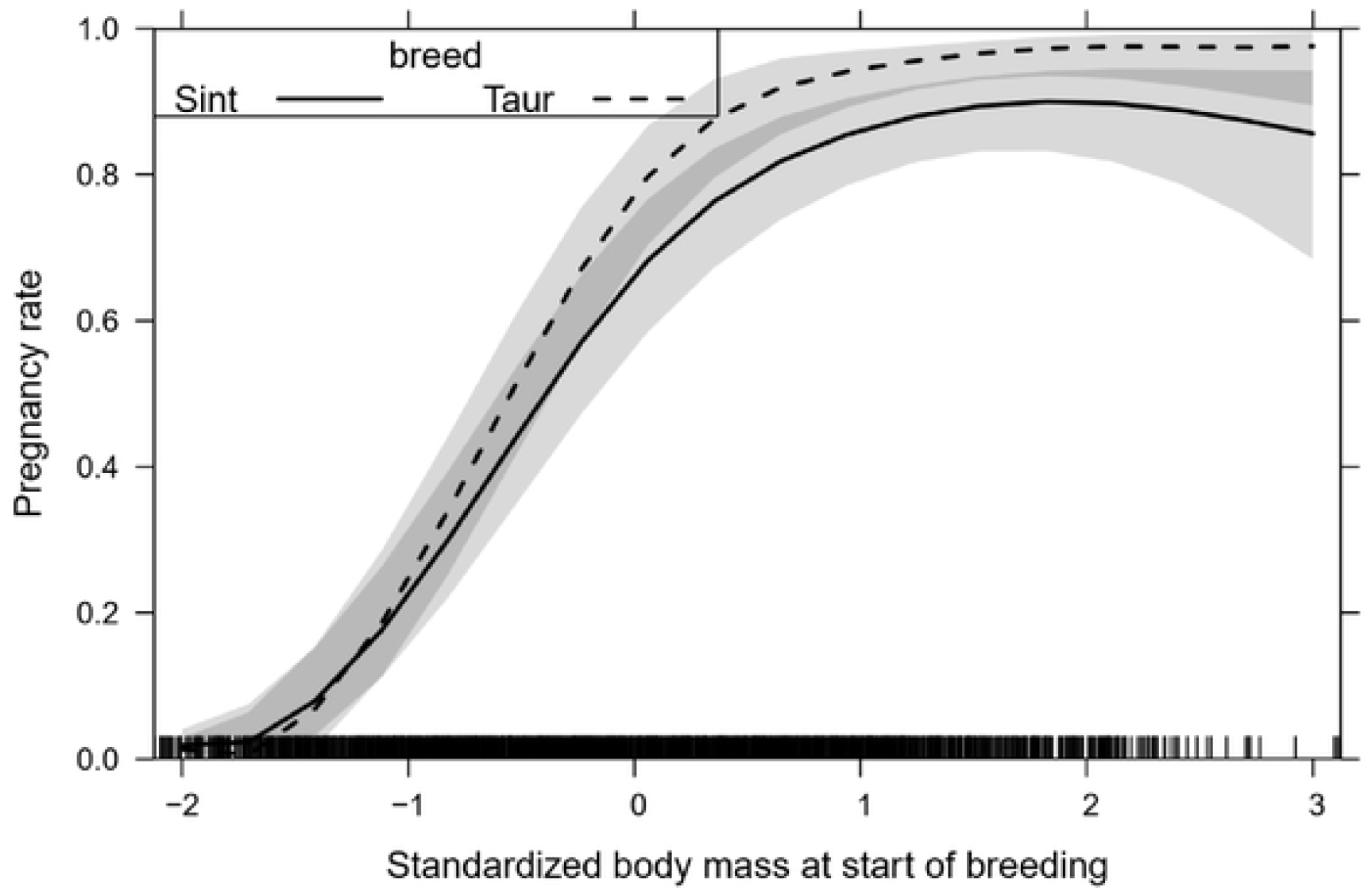
Interaction between body weight at the beginning of the breeding season and Breed to pregnancy rate of heifers and primiparous cows (Sint: crossbred, Taur: taurine females). Shaded strips represent 95 % confidence intervals for the expected value. Body mass average and standard deviation were 318 and 69 kg.

Pregnancy rate increased with increasing average daily gain during the breeding period. Averaging over other predictors, the model estimated that pregnancy rate increases from 59 % when animals lose 340 g per day to 79 % when they gain 890 g per day during the breeding season. When daily gain was at its average (0.264 ± 0.006 kg/day), pregnancy rate increased 1.7 % per 100 g of daily gain during the breeding period. When all covariates are at their average values for both categories, pregnancy rate was 25 percentage points higher for heifers than for primiparous cows (80 vs. 55 %).

### Effects of stocking rate not mediated by proximate factors

The addition of stocking rate as an explanatory factor resulted in the following model:

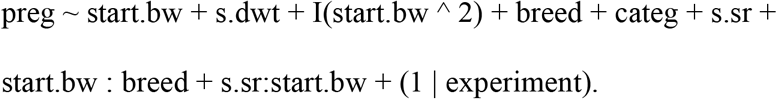

The contribution of stocking rate to explain variation in pregnancy rate was evaluated by entering stocking rate and its interactions last into the model and using a type II Wald test. Stocking rate and its interaction with body weight at the start of the breeding season contributed 4.7 % of the total sum of squares of the model and had direct effects on pregnancy rate that were significant even after controlling for the potential indirect effects through proximate variables such as body weight and weight change during breeding (Table 3). Stocking rate exhibited a significant interaction with weight at the start of the breeding season by which the effect of stocking rate was small for body weight below the 318 kg average and negative for heavier animals (Fig 3).

**Fig. 3.**
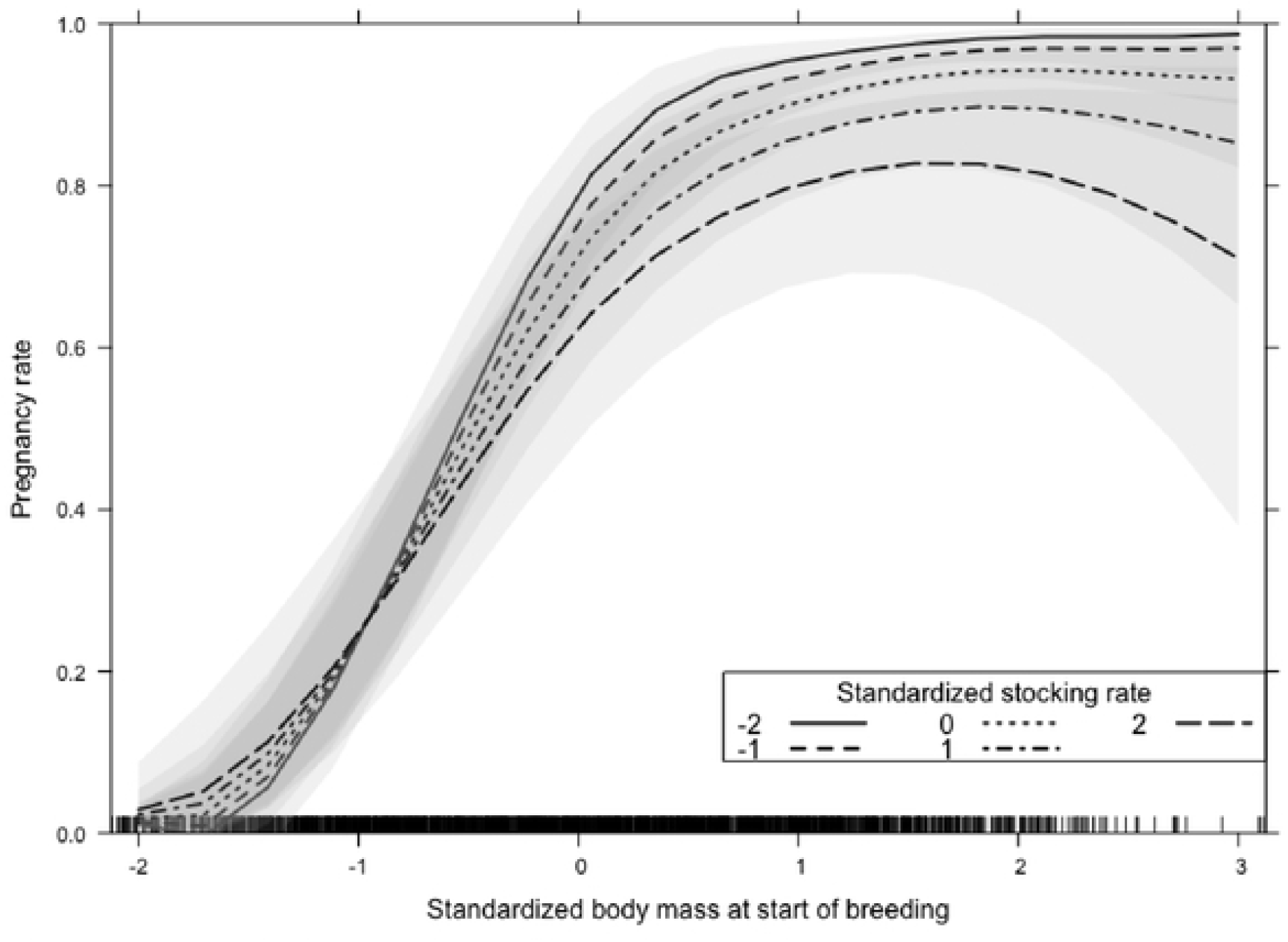
Interaction between body weight at the beginning of the breeding season and stocking rate on pregnancy rate of heifers and primiparous cows. Shaded areas are 95% confidence intervals. Tickmarks above the horizontal axis represent observations.

**Table 3.**
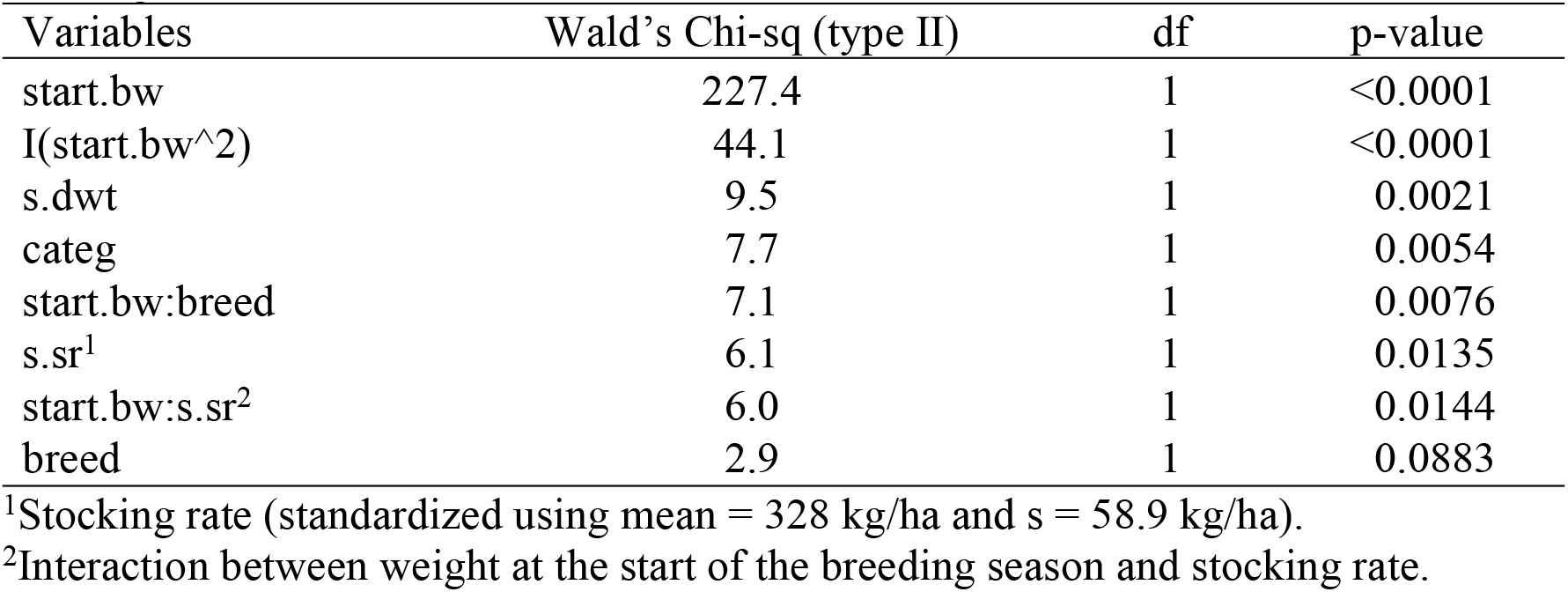
Analysis of variance of the pregnancy rate model with proximate factors and stocking rate

| Variables | Wald's Chi-sq (type II) | df | p-value |
| --- | --- | --- | --- |
| start.bw | 227.4 | 1 | <0.0001 |
| I(start.bw^2) | 44.1 | 1 | <0.0001 |
| s.dwt | 9.5 | 1 | 0.0021 |
| categ | 7.7 | 1 | 0.0054 |
| start.bw:breed | 7.1 | 1 | 0.0076 |
| s.sr<sup>1</sup> | 6.1 | 1 | 0.0135 |
| start.bw:s.sr<sup>2</sup> | 6.0 | 1 | 0.0144 |
| breed | 2.9 | 1 | 0.0883 |

<sup>1</sup>Stocking rate (standardized using mean = 328 kg/ha and s = 58.9 kg/ha).
<sup>2</sup>Interaction between weight at the start of the breeding season and stocking rate.

Although stocking rate did have effects on pregnancy rate beyond those through proximate causal variables, the effects of other variables did not change much by the incorporation of stocking rate. The most important factor affecting pregnancy rate when stocking rate was added in the model continued to be body weight at the beginning of the breeding season, accounting for 84.4 % of the sum of squares explained by the generalized mixed model (Table 3). The second largest contribution to the sum of squares of the model was due to average daily gain during the breeding season, which accounted for 4.3 % of the explained variation (Table 3). The response to daily gain was similar to that in the model without stocking rate; pregnancy rate increased 1.5 % per 100 g of daily gain when daily gain was near its average. The third largest contribution to the model’s sum of squares was due to animal category, accounting for 3.0 % of the explained variation. Pregnancy rate was greater in heifers than in primiparous cows (81 vs. 56 %, p = 0.0054).

### Body weight at the beginning of the breeding season

Because supplementation did not have detectable effects on body weight at the start of the breeding season, the final model was

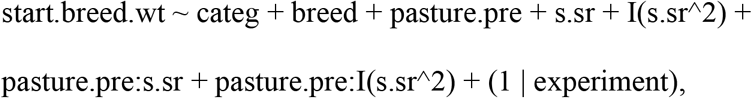

where *start.breed.wt* is body weight in kg at the start of the breeding season and *pasture.pre* is the type of pasture grazed prior to the breeding season. Thirty nine percent of the variation in initial body weight was explained by the fixed effects of the model, and an additional 41 % was explained by variation among experiments (random effects variation due to differences between experiments in variables not measured). Pasture type accounted for 60 % of the sum of squares explained by fixed effects, whereas an additional 27.9 % was explained by stocking rate and its interaction with pasture type. Body weight declined quadratically with increasing stocking rate in natural pastures, but it was not affected by the range of stocking rates studied in cultivated or improved pastures (Fig 4). At the average stocking rate of 328 kg/ha, initial body weight of animals grazing cultivated and improved pastures was 15 kg greater than that of animals grazing natural pastures, and this difference increased to 44 kg when stocking rate increased to 388 kg/ha.

**Fig. 4.**
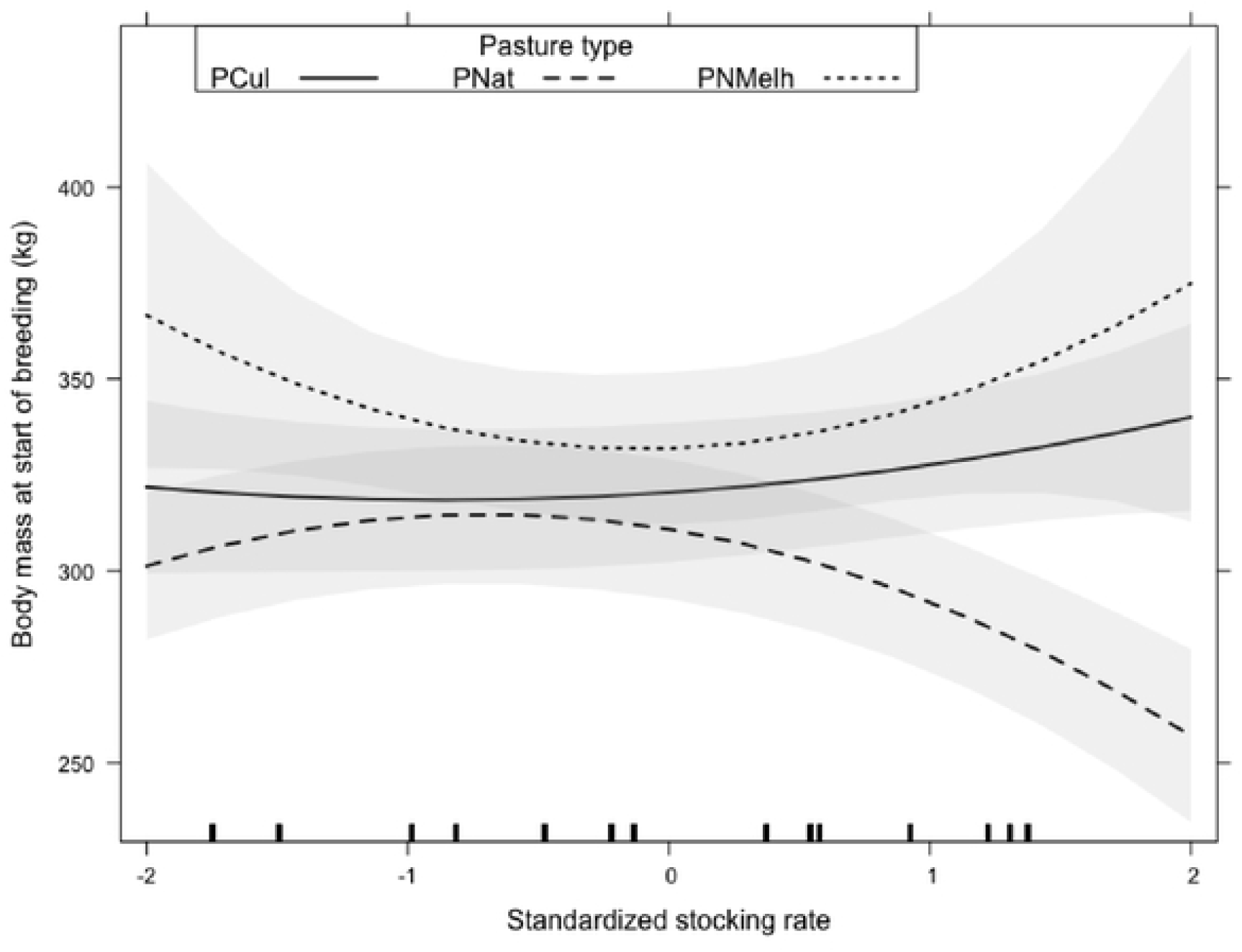
Effects of stocking rate and type of pasture on body weight at the beginning of the breeding season of heifers and primiparous cows. Efects of stocking rate were not significant for cultivated (PCult) and improved (PNMelh) pastures, but there was a significant quadratic effect in native grasslands (PNat).

Breed type and category had effects on initial weight that were independent of stocking rate. Ninety five percent confidence intervals for weight at the start of the breeding season were 303 to 338 and 295 to 330 kg for crossbred and taurine types.

Confidence intervals for heifers and primiparous cows were 263 to 313 and 332 to 380 kg.

This reflects the fact that multiple factors determine pregnancy rate, many of which are not closely related to body weight, animal category or breed. Other factors explored in some experiments, such as weaning method and use of artificial insemination, were considered and did not have detectable effects in preliminary models for pregnancy rate. However, the different weaning methods and use of artificial insemination were not represented across a good range of values in the other factors.

### Change in body weight during the breeding season

The final model for average daily gain during the breeding season was

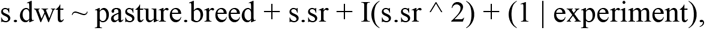

where *pasture.breed* is the type of pasture grazed during the breeding season.

Stocking rate and type of pasture grazed accounted for equal parts of the total variarion explained by the fixed effects of the model. Average daily gain during the breeding season decreased quadratically with increasing stocking rate (Fig 5).

**Fig. 5.**
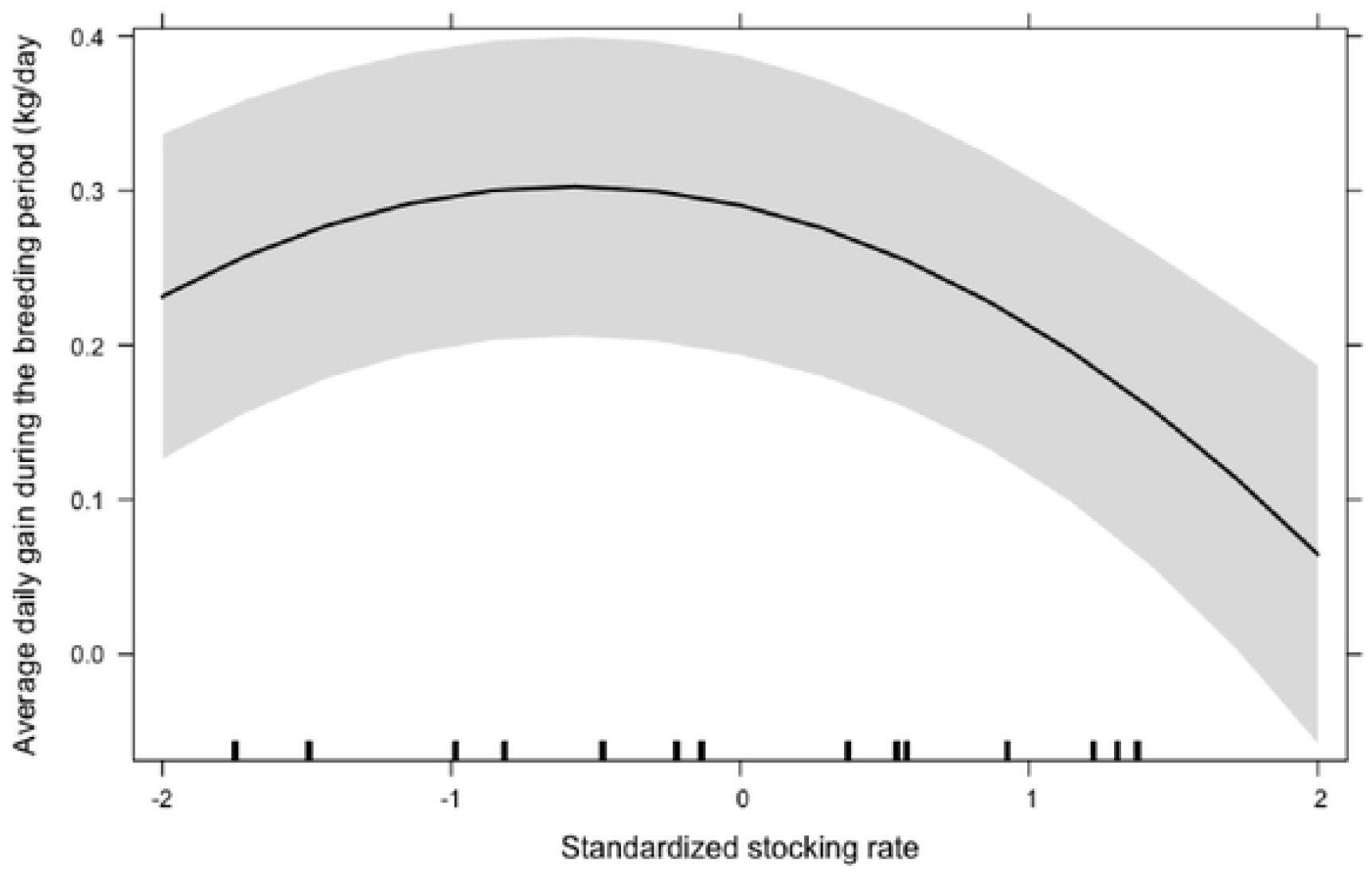
Average daily gain during the breeding season and stocking rate of heifers and primiparous cows. Shaded strip shows the 95 % confidence band.

## Discussion

This study is unique because it integrated data from thousands of individual animals from multiple sites and experiments and because it established quantitative relationships between reproductive performance and stocking rate. Although stocking rate is recognized as one of the most important factors determining productivity of grazing systems (Hunt, Mclvor, Grice & Bray 2014), studies relating grazing animal performance to stocking rate are rare. A Web of Science search performed on 04 Jan 2020 with the terms “beef catle” AND “pregnancy rate” AND “stocking rate” yielded nine articles, only one of which (Renquist, Oltjen, Sainz, Connor & Calvert 2005) presented original data on the effects of stocking rate on pregnancy rates. Most of the studies where stocking rate is considered as one of the explanatory variables for animal performance include few levels of stocking rate that explore a very limited range. We surmise that as a consequence of the limited range explored and the inherent high variability of herd-level studies, many studies failed to detect effects of stocking rate. The present study included 16 levels of stocking rate ranging from 0.5 to 1.2 head/ha or 200 to 464 kg/ha, which allowed quantification of response curves.

### Effects of proximate causal factors on pregnancy rate

The effects of body weight at the beginning of the breeding season, animal category and changes in body weight during the breeding season were quantified and yielded response curves with low variance. For example, the pseudo coefficient of variation (CI half width/(2 expected value) of pregnancy rate for taurine cows at average weight at the beginning of the breeding season was 6.5%. The most important causal factor influencing pregnancy rate was body weight at the beginning of the breeding season, which interacted with breed to determine that at high initial weights, taurine females had higher pregnancy rate than crossbred females (Fig 2). Pregnancy rate is influenced by nutrition, because it directly affects the reproductive physiology in beef cows [6], mainly in periods of higher requirements like pre and postpartum. If nutrition is inadequate, body reserves become depleted and body condition declines (Diskin & Kenny 2016), resulting in low ovulation rate. Females with adequate metabolic status and high body weight have high levels of glucose, insulin and growth factor I (IGF-I) (Yelich *et al.* 1995; Santos & Amstalden 1998), potentiating the effect of gonadotrophins (LH and FSH) (Spicer & Echternkamp 1995) and promoting ovulation (Sirois & Fortune 1988). Pre and postpartum periods coincide with low availability of nutrients in natural grasslands, which are characterized by variation in composition, structure and, seasonality of production and quality (Mezzalira *et al.* 2014; Neves *et al.* 2009).

Animals with smaller frame reach physiological maturity earlier, at a lower weight and with greater fat content than larger animals (Owens, Dubeski & Hanson 1993). When growth rate decreases and the process of fat deposition begins, larger animals are still in the growth phase (Mckiernan 2005). In addition, the higher pregnancy rate observed in the taurine females can be explained by the greater selection for precocity carried out in the herds from which these females proceed (Diskin & Kenny 2016). When heifers reach puberty and mate earlier the biological efficiency of the herd is improved for as long as the early mating does not compromise full development. These characteristics become more important as production systems become more intensive and competitive. Reducing the age at first conception alters the structure of the herd and shortens the interval between generations, thus decreasing the participation of unproductive animals in the composition of the herd (Pötter, Lobato & Mielitz Neto 1998; Beretta, Lobato & Mielitz Neto 2001).

Weight gain during the breeding season is clearly important for cows to become pregnant. Greater weight gain in this period indicates that forage is less limiting, and that sufficient quantity and quality of food intake is obtained to support ovarian activity (Vieira, Lobato, Torres Junior, Cezar & Correa 2005). According to Carter & Cox (1973), there is greater biological efficacy in females that have their first calf at about two, rather than three or more years of age. Adequate weights at the beginning of the breeding season are decisive for a high conception rate (Wiltbank, Rowden, Ingalls, Geegoey & Koch 1962).

The lower pregnancy rate of primiparous cows than heifers may be related to the stress of calving and the combined effects of growth and first lactation requirements of primiparous cows. Low reproductive success has been documented for primiparous animals when they are subjected to periods of pre or postpartum feeding restriction (Spitzer, Morrison & Wettemann 1995). The negative direct effect of primiparous condition on pregnancy rate appeared with the inclusion of initial body weight, but primiparous condition had an indirect positive effect on pregnancy rate relative to heifers through the fact that they were heavier than heifers at the beginning of the mating period (Fig 6).

**Fig. 6.**
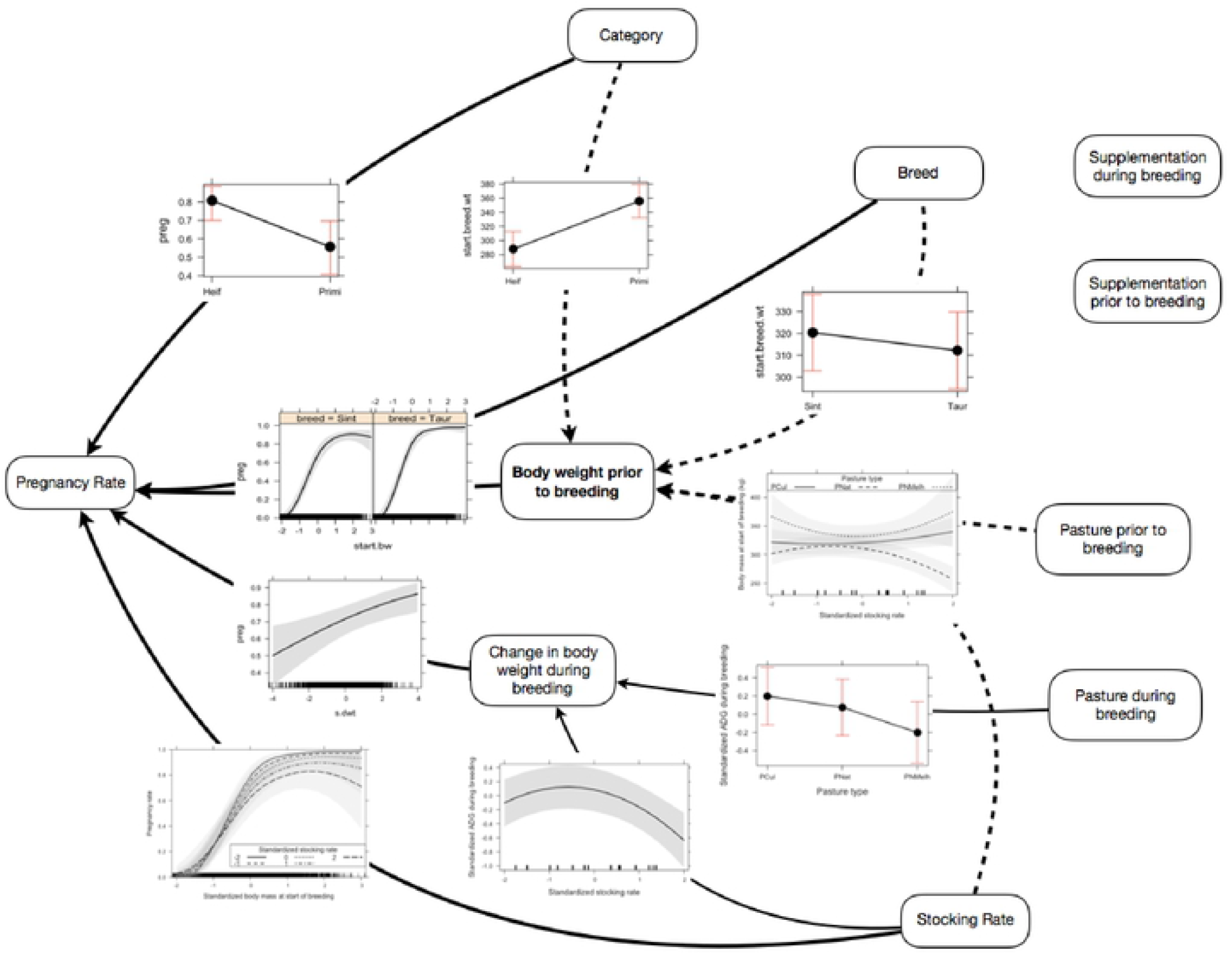
Schematic representation of modeling results showing effects of animal factors (category and breed type), foraging environment (pasture type, stocking rate and supplementation) on body weight and reproductive performance of beef cattle. In the broader lines are observed direct effects on the pregnancy rate. In the thinner lines are observed indirect effects of body weight prior to breeding and in fine and dotted lines, indirect effects of the change in body weight during breeding.

Stocking rate interacted with body weight at the beginning of the breeding season, whereby higher stocking rates were associated with lower pregnancy rate only in the high range of body weight (Fig 3). Lower stocking rates allow greater development of the animal, due to the higher forage accumulation, making it possible for females to have more food available (Euclides & Euclides Filho 1998). Stocking rate is a primary management variable in grazing systems because it modulates the interactions between animals and pasture (Bransby & Maclaurin 2000; Carvalho & Batello 2009). As stocking rate increases, herbage allowance decreases and can reach levels where intake per animal is too low for production, but intake per unit area surpasses the ability of pastures to produce and recover. Individual animal performance decreases as stocking rate increases, because the daily intake is constrained by limiting sward structure at low herbage allowance (Da Trindade *et al*. 2014), but production per unit area increases and then decreases with increasing stocking rate (Mezzalira *et al*. 2012; Neves *et al*. 2009; Petersen, Lucas & Mott 1965). High stocking rates pre and post partum make it difficult for cows to recover good body condition after calving, compromising the reproductive performance of the cow and the productivity in subsequent seasons and reproductive years (Osoro 1989). Therefore, stocking decisions must be informed by curves that relate individual performance to stocking rate like the ones provided in the present study. These curves are particularly important for the integration of biological and economic functions to determine optimal stocking rates.

As expected, heifers achieve greater pregnancy rates when mantained in good nutritional conditions. Inadequate management practices, such as excessive stocking rate and lack of custom management for certain animal categories have led to generally low indices of productivity in the region. However, there are possibilities for reducing the age of slaughter and the age at first breeding, which may allow the improvement of productive and reproductive indices (Pötter, Lobato & Mielitz Neto 2000; Beretta, Lobato & Mielitz Neto 2002). Because natural pastures of the region are dominated by warm season grasses with low productivity and quality in the cold season, grazing of cultivated cool-season pastures significantly increases indicators of reproductive performance, beef yield and economic results (Lupatini & Neumann 2002).

In agreement with (Rovira 1974), we observed that when heifers and primiparous cows are mantained in optimal conditions of grazing and nutrition, that is, mantained in high quality pastures, with sufficient body weight and intermediate stocking rates, they achieve near maximal reproductive success. Sufficient nutrition allows early breeding, which increases the overall efficiency of production for the herd.

### Effects of stocking rate not mediated by proximate factors

Our results show that stocking rate has an effect on pregnancy rate that is not explained by any of the other variables considered. Even after controlling for effects of starting body weight and weight change during the breeding season, when body weight at the beginning of the breeding season is greater than average, pregnancy rate declines with increasing stocking rate. This effect of stocking rate appears to be restricted to the range of starting body weight where pregnancy rate no longer responds to body weight. This further suggests that stocking rate had an effect that was not mediated by the observed effects of stocking rate on body weigh at the start of the breeding period and weight change during breeding (Fig 6).

Effects of stocking rate on pregnancy rate that are not related to nutritional condition, as reflected in body weight and daily gain, might be related to animal health and associated management variables. Higher stocking rates may result in greater load of external and internal parasites (Gasser 2013). Tick infestation is common in this region, and ticks frequently carry Babesia (Bilhasi *et al*. 2014). When grazing at higher stocking rates, animals are forced to graze closer to the soil and increase the rate of ingestion of parasite helminth larvae.

### Body weight at the start of breeding season and changes in body weight during the breeding season

Our results agree with the conventional wisdom that stocking rate is one of the most important factors in grazing management. Stocking rate interacting with type of pasture before mating was the most important factor influencing body weight at the beginning of the breeding season (Fig 4) and it was the most important factor affecting the changes in body weight during the breeding season (Fig 5). The lowest weights at the beginning of the breeding season observed in this study are close to the minimum weight recommended for the first breeding season (50 - 57 % of adult weight) to avoid impairment of life-long reproductive performance (Funston & Deutscher 2004; Roberts, Geary, Grings, Waterman & MacNeil 2009). Body weight of heifers and cows grazing cultivated or improved pastures was high and did not respond to stocking rate, presumably because the level of feeding and pasture production was sufficient to provide enough nutrition at all stocking rates studies. On the other hand, we observed a typical response of declining in body weight and average daily gain during the breeding season for animals grazing native pastures where forage amount and quality are lower than in cultivated and improved pastures. Lower stocking rates allow animals to select better-quality diets, while higher stocking rates reduce vegetation abundance, constraining daily intake (Da Trindade et al., 2016).

## Conclusions

This synthesis of a large number of experiments conducted over decades in the Pampas region confirms the importance of body weight at the start of the breeding season to achieve high pregnancy rates in cattle. Body weight at the beginning of the breeding season is an easily measurable variable and can be used herd reproductive management. Stocking rate had a negative effect on pregnancy rate both through its negative effects on initial body weight and weight change during the breeding season, and its direct negative effects on pregnancy rates when body weight was not limiting. Heifers tended to have lower body weight than primiparous cows, but after correction for body weight, they had greater pregnancy rates than primiparous cows, most likely due to the fact that primiparous cows were simultaneously lactating and growing. Response curves derived from out study can be used to optimize stocking rates under various economic conditions and to guide policies to improve the efficiency of reproductive livestock herds under free-grazing conditions.

